# Paradoxical Th1 activation and CTLA-4 regulation is beneficial during latent cryptococcosis

**DOI:** 10.64898/2026.07.02.736061

**Authors:** M Ding, J Drnevich, JM Yoder, EV Dang, K Nielsen

## Abstract

*Cryptococcus neoformans* is the predominant causative agent of cryptococcal meningitis in immunocompromised individuals. Conversely in immunocompetent individuals, *C. neoformans* establishes a latent pulmonary infection characterized by a paucity of clinical symptoms. Using a mouse inhalation model of latent *C. neoformans* infection, we previously showed that CD4 T-cells are necessary for preventing fungal proliferation in the lungs. In the current study, we performed single cell RNA sequencing (scRNAseq) and found that the CD4 T-cell response was both highly heterogenous and dichotomous during pulmonary *C. neoformans* infection, with concomitant expression of genes related to Th1 polarization (*Tbx21*, *Ifng*) and immune regulation (*Ctla4*). First, we demonstrated that cells with Th1-like phenotypes are necessary and sufficient to control latent infection via adoptive transfer of T-bet positive cells into infection-matched CD4-depleted recipient mice. Second, scRNAseq analysis revealed the subpopulation of effector CD4 T-cells that co-expressed *Ctla4* and *Gata3* was significantly higher than a subpopulation that co-expressed *Ctla4* and *Tbx21*. Furthermore, our data suggested that CTLA-4 upregulation is beneficial against *C. neoformans* infection, as CTLA-4 blockade promoted fungal proliferation. Thus, we propose a model wherein Th1 control of latent *C. neoformans* infection is supported by CTLA-4 suppression of detrimental Th2 activation.

**IMPORTANCE:** CD4 T-cells are a critical part of the adaptive immune response in many infections. Upon inhalation of the fungal pathogen *Cryptococcus neoformans*, CD4 T-cells are necessary to control the initial latent infection and prevent disease. In immunocompromised individuals that lack CD4 T-cells, the initial infection in the lungs can spread to the brain and cause lethal meningitis. In this study, we show that the CD4 T-cells present during latent *C. neoformans* infection are heterogeneous with expression of diverse polarization and immune regulation markers that function in concert to prevent disease. Our studies highlight the previously unappreciated diversity and complex regulation of the CD4 T-cell response that is required to prevent disease in this important fungal pathogen.

## INTRODUCTION

*Cryptococcus neoformans* is an opportunistic fungal pathogen that causes life-threatening cryptococcal meningitis in immunocompromised individuals. It is the second most common HIV/AIDS-associated infection world-wide, with an estimated mean global cryptococcal antigenemia prevalence of 4.4% among people with CD4 counts less than 200 cell/μL (1, 2). Following the inhalation of spores or desiccated yeast cells, *C. neoformans* establishes an initial pulmonary infection that is contained within granulomas (3–5). In healthy individuals, *C. neoformans* is able to persist within the lungs in the absence of any clinical symptoms (6). However, this latent infection can disseminate from the lungs and progress to fatal cryptococcosis in immunocompromised individuals (6).

Characterization of the host immune response to latent *C. neoformans* infection is limited. The majority of our knowledge about the host response to *C. neoformans* infection is derived from mouse studies using highly virulent *C. neoformans* strains that establish lethal infections, such as KN99α (7–9) and H99 strains (10, 11). We recently developed a mouse inhalation model of latent *C. neoformans* infection, utilizing the fully-virulent *C. neoformans* clinical isolate UgCl223 (12). With this mouse model, we demonstrated that depletion of the CD4 T-cells resulted in disseminated lethal meningitis and lung Tbet+ Th1 cells appeared to be the predominant canonical CD4 T-cell subset (12). Surprisingly, the proportion of activated pulmonary Th1 cells expressing IFNγ upon PMA/ionomycin restimulation is significantly decreased during latent infection compared to that of uninfected controls (12). These findings suggested that while Th1 cells are important in the host adaptive immune response to latent *C. neoformans* infection, they may be unable to promote complete clearance of the infection, likely due in part to diminished capacity to produce IFNγ. However, it is unknown what other components of the pulmonary CD4 response contribute towards preventing fungal proliferation and mortality during latent *C. neoformans* infection.

In the current study, we utilized scRNAseq to characterize the heterogeneity of the effector CD4 T-cell response to latent and lethal *C. neoformans* infection. Our analysis revealed a complex network of effector CD4 T-cell subsets with seemingly antagonistic qualities. To explore these paradoxical immune responses, we parsed out the interplay between the pro-inflammatory Th1 response and a regulatory CTLA-4 response using adoptive transfer and antibody blockage studies to elucidate the complex role these diverse CD4 T-cells play in control of latent *C. neoformans* infection.

## RESULTS

### Heterogeneity of CD4 T-cell responses at the single-cell level

We previously showed that CD4 T-cells are necessary for controlling latent *C. neoformans* infection, with depletion of CD4 T-cells resulting in a significant increase in lung and brain fungal burdens in infected mice and subsequent mortality (12). Our findings also demonstrated that Tbet-expressing Th1 cells were prevalent during latent *C. neoformans* infection (12). This contrasts with lethal *C. neoformans* infections where detrimental Th2 cells predominate (9). Surprisingly, we identified Tbet+ and Gata3+ Th1/Th2 hybrid cells within the lungs of latently infected mice that implied the host response could be more heterogenous than previously theorized (12). Thus, we utilized single cell RNA sequencing (scRNA-seq) analysis to define the heterogeneity of the host response to latent *C. neoformans* infection and resolve the individual subsets of major immune cell populations.

We initially performed a preliminary study to examine the total lung immune response to *C. neoformans* infection. CD45+ cells were isolated from lung homogenates of a mouse latently infected with UgCl223 at 14 days post-infection (DPI), a mouse latently infected with UgCl223 at 150 DPI, a mouse lethally infected with KN99α at 14 DPI, and an uninfected mouse. Using scRNAseq analysis, we classified the lung CD45+ cells into major immune cell populations using reference-based annotation (**Supplemental Figure 1A-B**). T-cells were the only population in which we identified a visibly dramatic shift in cluster size between the different experimental conditions (**Supplemental Figure 1C**). ScRNAseq analysis of lung CD45+ cells proved very useful in understanding the overall transcriptional heterogeneity of the pulmonary immune population during *C. neoformans* infection, but using bulk CD45+ lung cells limited our ability to resolve the CD4 T-cell subpopulations during the *C. neoformans* infection (**Supplementary Figure 1D**). Although differential expression analysis of the CD4 T-cells showed a significant change in some immunomodulatory genes (**Supplementary Figure 1E**), additional analysis was limited due to the low number of *Cd4* expressing cells present in the total leukocyte populations.

To enhance the detection of the CD4 T-cell populations, we designed a more focused scRNAseq experiment using CD4 T-cells isolated via negative selection from lung homogenates of mice latently infected with UgCl223 at 14 DPI, latently infected with UgCl223 at 180 days DPI, and lethally infected with KN99α at 14 days DPI. Following scRNAseq analysis, we identified 10 CD4 T-cell subsets (**Figure 1A**). Only Cluster 2 were effector T-cells based on their high expression of *Cd44*, a marker of T-cell activation (**Figure 1B**). The remaining CD4 T-cell subsets had significantly increased expression of the naïve CD4 T-cell genes *Ccr7*, *Sell*, and *Lef1* (13, 14) (**Figure 1C**). Although the top 10 expressed genes in the naïve CD4 T-cell subsets were unique (**Supplementary Figure 2**), many of the genes were not previously studied in the context of CD4 T-cells and this limited our ability to characterize the naïve CD4 T-cell subsets (**Figure 1D**). Of note, we did observe expression of genes involved in T-cell activation, such as *Camk1d* and *Ifit3*/*Ifit1* (**Supplementary Figure 2**) (15, 16), as well as a noticeable change across experimental conditions in naïve cells expressing heat shock genes (cluster 5) and the effector cell populations (cluster 2) (**Figure 1D**).

**Figure 1:**
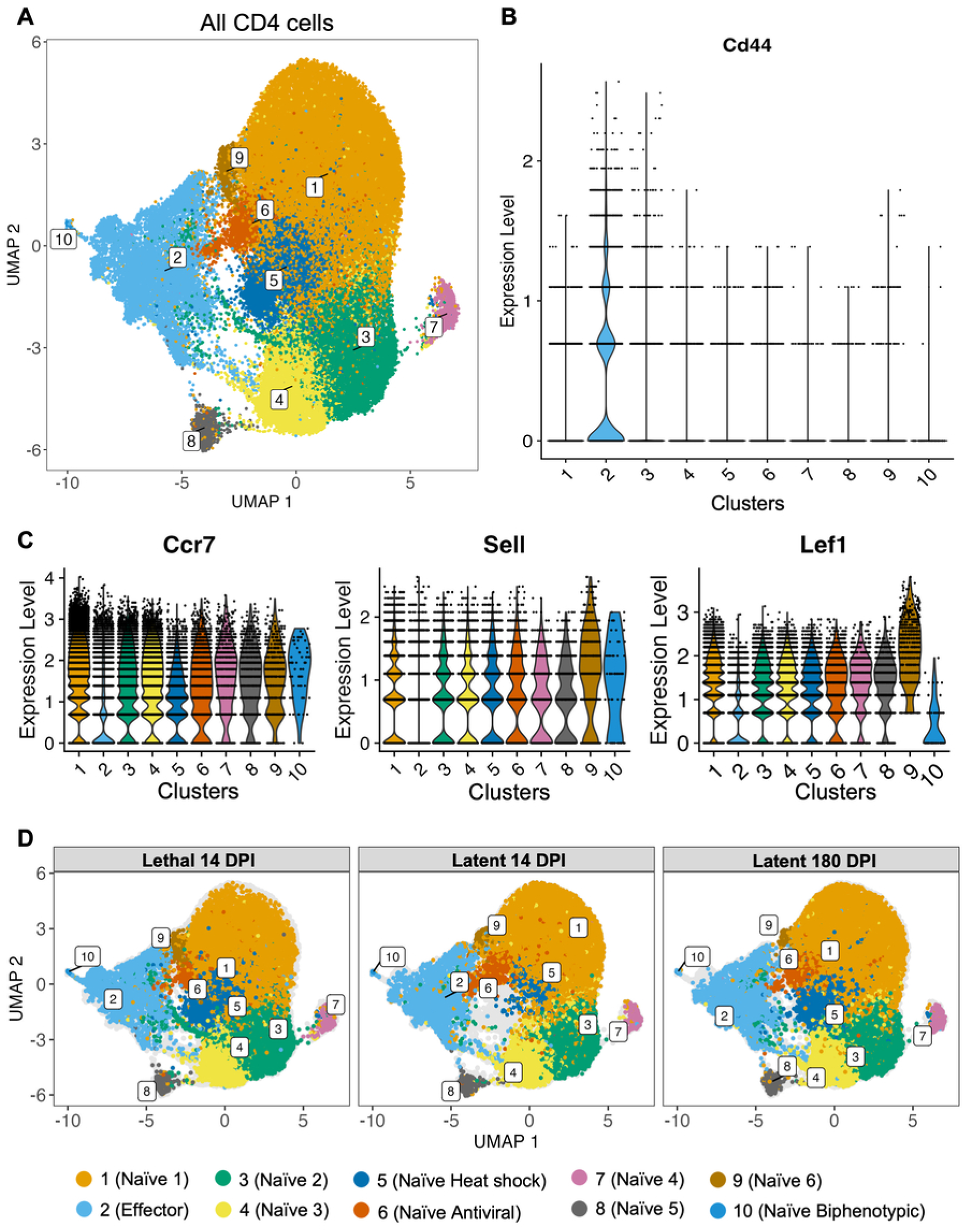
Single-cell RNA sequencing analysis of pulmonary CD4 T-cells during *C. neoformans* infection. C57BL/6J mice were infected with *C. neoformans* KN99*α* at 14 days post-infection (Lethal 14 DPI), UgCl223 at 14 days post-infection (Latent 14 DPI), and UgCl223 at 180 days post-infection (Latent 180 DPI). CD4+ cells were isolated from lung homogenates via negative selection and processed for single-cell RNA sequencing analysis via the 10x Genomics platform. (A) UMAP projection showing Seurat clustering of CD4 T-cells. (B) Violin plot of *Cd44* expression (marker of T-cell activation) across all Seurat clusters. (C) Violin plots of *Ccr7*, *Sell*, and *Lef1* expression (markers of naïve T-cells) across all Seurat clusters. (D) Individual UMAPs of the 3 experimental conditions were labelled based on the inferred subtype using the top 10 expressed genes in each Seurat cluster (**Supplementary Figure 2**).

Based on these findings, we narrowed our subsequent analyses to the *Cd44*-expressing effector CD4 T-cell subset in Cluster 2. Additional clustering analysis of the *Cd44*-expressing cells (cluster 2) revealed 12 effector CD4 T-cell subclusters (**Figure 2A**). The top 10 expressed genes with known functions in the effector CD4 T-cell subclusters were used for annotation (**Figure 2B, Supplementary Figure 3**). Subcluster 1 was defined as lymph node-derived based on high expression of the lymph node egress genes *S1pr1* and *Klf2* (17). Subcluster 2 was defined as transitional based on high expression of chemokine receptor gene *Cxcr6* involved in mediating inflammation (18), and immune polarization genes *Rbpj*, *Ccr8*, and *Bhlhe40* (19–24). Subcluster 3 was defined as regulatory T-cells (Treg 1) based on high *Foxp3* expression (25). Subcluster 4 was defined as central memory cells based on high expression of chemokine receptor gene *Ccr7* (26, 27). Subcluster 5 was defined as an antifungal effector cell cluster based on high expression of the *Myo1f* gene, which was previously implicated in antifungal immunity against *Candida albicans* and *Aspergillus fumigatus* (*28, 29*). Subcluster 6 was defined as a second regulatory T-cell subcluster (Treg 2) with high co-expression of *Foxp3* and *Areg* (30), along with expression of the immune exhaustion gene *Ctla4* (31). Subcluster 7 was defined as cytotoxic gamma-delta T-cells based on high expression of the cytotoxic genes *Nkg7* and *Eomes* (32, 33), and high expression of the gamma T-cell receptor gene *Trgv2* (34). Subcluster 8 was defined as a cluster of pro-inflammatory effector cells based on high expression of the proinflammatory genes *Ahnak* and *Rnf19b* (35–38). Subcluster 9 was defined as exhausted CD4 T-cells based on high expression of exhaustion associated genes *Tox2*, *Sostdc1*, and *Lag3* (39–41). Subcluster 10 was defined as Th2-like gamma-delta T-cells based on high expression of the type-2 associated gene *Arg1* (42), and the delta and gamma T-cell receptor genes *Trdv4* and *Tcrg-C4*, respectively (34). Subcluster 11 was defined as heat shock-related CD4 T-cells based on high expression of the heat shock genes *Hspa1a* and *Hspa1b*. Finally, subcluster 12 was defined as unconventional gamma-delta T-cells based on high expression of the gamma-delta transcriptional regulator gene *Sox13* (43), gamma T-cell receptor gene *Tcrg-V4* (34), innate-like unconventional T-cell development regulator gene *Zbtb16* (*44*), and pro-inflammatory IL-23 receptor gene *Il23r* (45, 46) (**Figure 2C**, **Supplementary Figure 3**).

**Figure 2:**
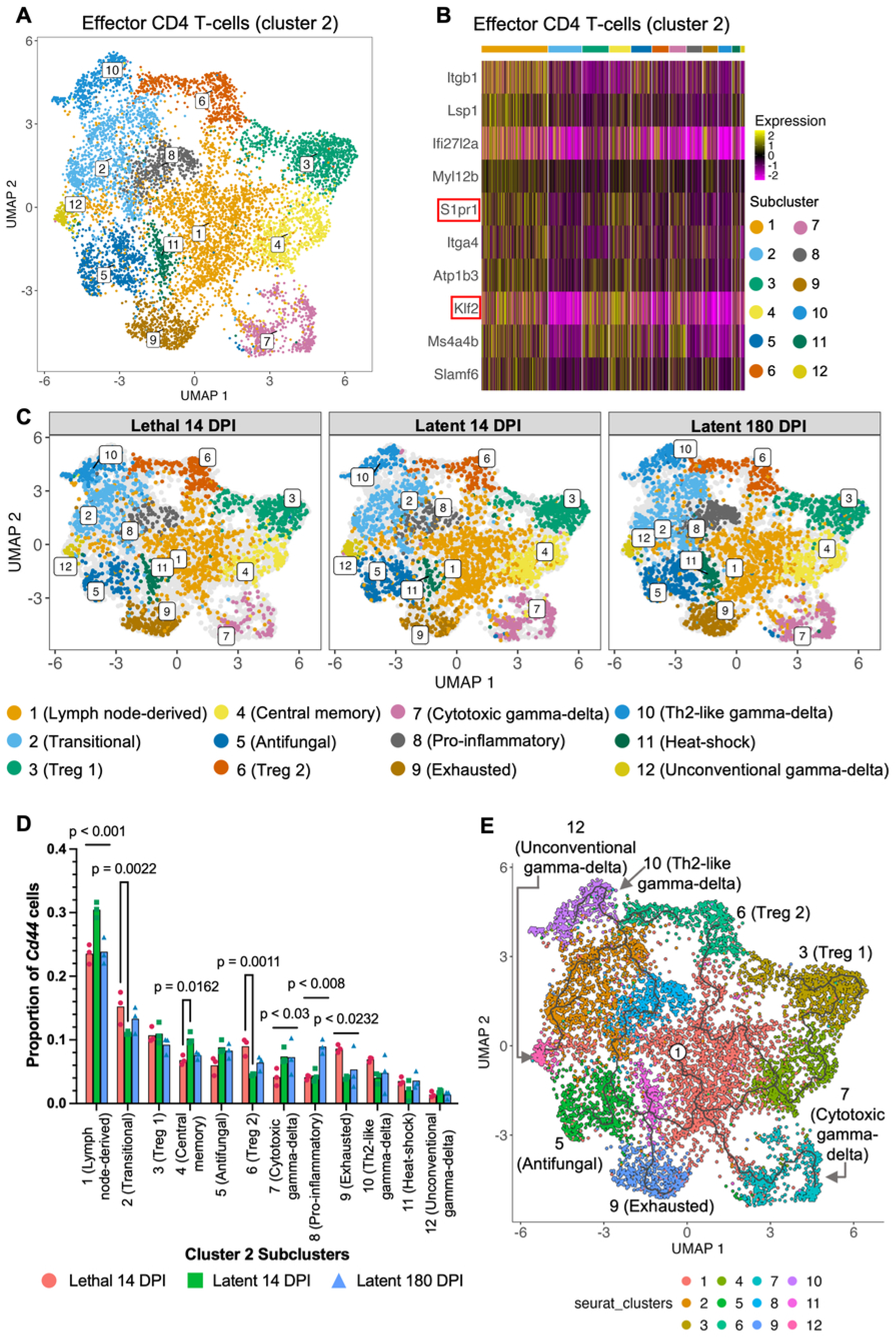
Single-cell RNA sequencing analysis of pulmonary effector CD4 T-cells during *C. neoformans* infection. (A) UMAP projection of effector CD4 T-cells (cluster 2) by Seurat clusters. (B) Heatmap of top 10 genes expressed by the effector CD4 T-cell cluster (cluster 2), with yellow to purple intensity indicating the mean expression of each gene and the cluster number indicated by the colored bar. (C) UMAP split according to experimental condition: *C. neoformans* KN99*α* at 14 days post-infection (Lethal 14 DPI), UgCl223 at 14 days post-infection (Latent 14 DPI), and UgCl223 at 180 days post-infection (Latent 180 DPI), and labelled according to inferred subtype based on the top 10 expressed markers in the Seurat cluster (**Supplementary Figure 3**). (D) Proportion of pulmonary effector CD4 T-cells in the Seurat clusters across the experimental conditions: Lethal 14 DPI, Latent 14 DPI, Latent 180 DPI. For each experimental condition, the proportion was calculated by dividing the number of cells in each effector CD4 T-cell subcluster by the total number of effector CD4 T-cells. Significance was determined by two-way ANOVA with Bonferroni correction. (E) Branched pseudotime trajectory of effector CD4 T-cell activation/differentiation inferred by Monocle 3 with the lymph-node subcluster as the root. Each cell was colored based on Seurat clustering and the end trajectories of the branch points were labelled.

When examining differences in the proportion of cells in the effector CD4 subclusters, we noted a few significant differences. Specifically, the proportion of cytotoxic gamma-delta T-cells (subcluster 7) was significantly decreased and the proportion of exhausted CD4 cells (subcluster 9) was significantly increased in the lethal infection when compared to both latent infection timepoints (**Figure 2D**). In addition, the proportion of pro-inflammatory effector CD4 T-cells (subcluster 8) was increased during the 180 DPI latent infection, and the proportion of lymph node-derived CD4 T-cells was higher during 14 DPI latent infection, compared to the other two experimental conditions, respectively (**Figure 2D**). Aside from these quantitative differences, the overall UMAP projection amongst the three experimental conditions was similar in appearance (**Figure 2C**).

The progression of effector T-cell activation and differentiation was inferred using pseudotime trajectory analysis of the subclusters. We selected the lymph node-derived subcluster (subcluster 1) as the root node. The expression of the lymph node egress genes *S1pr1* and *Klf2* (17) suggested that they were new arrivals into the lung environment. In addition, these cells also expressed the T-cell proliferation regulatory gene *Ms4a4b* (47, 48) and the T-cell activation gene *Slamf6* (49, 50) (**Figure 2B**). The pseudotime analysis revealed multiple branch points, each with distinct trajectories (**Figure 2E**). Notably, the majority of trajectories ended at the following clusters: Treg 1, antifungal effector cells, Treg 2, cytotoxic gamma-delta T-cells, exhausted CD4 T-cells, Th2-like gamma-delta T-cells, and unconventional gamma-delta T-cells (**Figure 2E**). Overall, the pseudotime analysis demonstrated that many of the effector CD4 subsets may be precursors or intermediates during CD4 polarization.

### Increased Type-1 polarization of effector CD4 T-cell responses during latent infection

Transcriptional differences between the effector cell subsets during the latent and lethal infections were analyzed via differential gene expression analysis. First, we compared gene expression between the lethal infection at 14 DPI and the latent infection at the same 14 DPI timepoint (**Figure 3A**). The differential expression analysis showed significant upregulation of the Th2-associated genes *Il4*, *Il5*, and *Il13*, and the type-2 associated gene *Arg1* in the lethal infection. This was coupled with significant upregulation of the immune suppression genes *Foxp3, Il10ra, Il10*, *Il2ra,* and *Nrp1* in the lethal infection. (**Figure 3A**). However, there was also significant upregulation of genes related to immune cell regulation and inhibition (*Lag3*, *Pdcd1*, *Ctla4*, *Tigit*) in the lethal infection compared to the latent infection (**Figure 3A**). Combined with the significant increase in the proportion of exhausted CD4 T-cells during lethal infection, these findings indicated that the lethal *C. neoformans* infection had a robust Th2-mediated CD4 T-cell response with increased immune suppression and inhibitory gene expression when compared to the latent infection.

**Figure 3:**
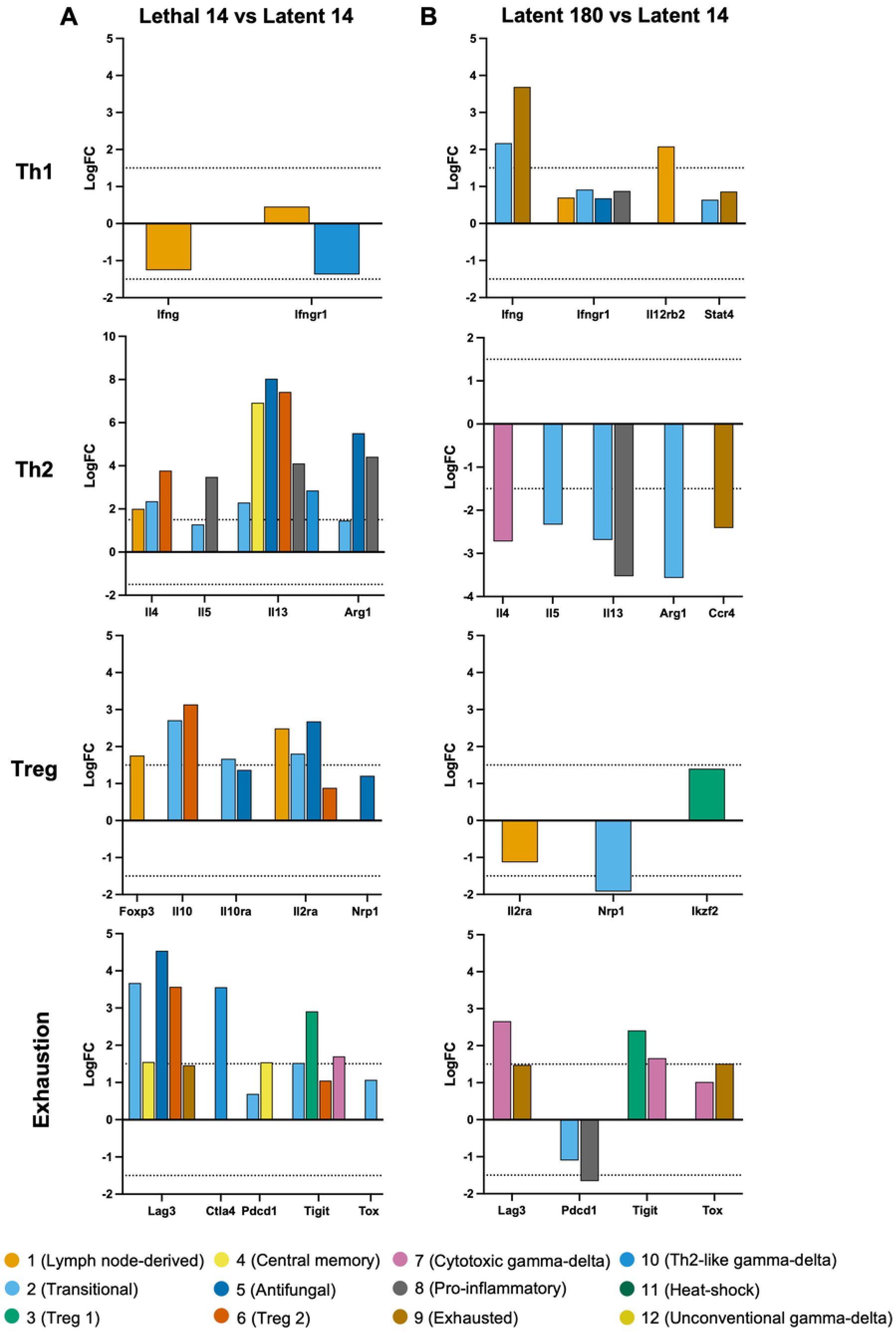
Th1-related genes are increased and Th2-related genes are decreased over the course of latent *C. neoformans* infection. Differential expression analysis of pulmonary effector CD4 T-cell subclusters 1-12 was performed between two experimental conditions: (A) *C. neoformans* KN99*α* at 14 days post-infection (Lethal 14 DPI) and UgCl223 at 14 days post-infection (Latent 14 DPI), and (B) *C. neoformans* UgCl223 at 180 days post-infection (Latent 180 DPI) and Latent 14 DPI. For each comparison, significant Th1-, Th2-, Treg-, and exhaustion-associated genes within each subcluster (with globally adjusted p-value < 0.05) were plotted according to logFC. Positive logFC values indicate higher expression in the first condition listed, whereas negative logFC values indicate higher expression in the second condition. The dashed horizontal line denotes biological significance at |logFC| ≥ 1.5.

Second, we analyzed gene expression differences between the early latent infection 14 DPI timepoint compared to the late latent infection timepoint at 180 DPI (**Figure 3B**). The early timepoint served as a useful control and represented an immune state where fungal burden is not yet optimally controlled, whereas the late timepoint was chosen to occur after the pulmonary granuloma has effectively contained and controlled the *C. neoformans* infection (51). This analysis revealed significantly increased expression of the Th1-associated cytokine gene *Ifng* (IFNγ) over time (**Figure 3B**). Interestingly, we also observed an increase in the immune inhibitory genes *Lag3* and *Tigit* in the cytotoxic gamma-delta (subcluster 7) and Treg 1 (subcluster 3) effector CD4 T-cell populations, respectively, in the late stage of the latent infection (**Figure 3B**). However, we also observed a decrease in the immune inhibitory gene *Pdcd1* in the pro-inflammatory (subcluster 8) effector CD4 T-cell population (**Figure 3B**). Finally, there was a significant decrease in the Th17-associated gene *Il17a* (**Supplementary Figure 4**) and the Th2-associated genes *Il4*, *Il5*, *Il13*, *Arg1*, and *Ccr4* (**Figure 3B**) in the late stage of the latent infection.

Indeed, the proportion of all effector CD4 T-cells expressing the Th1 transcription factor gene *Tbx21* was significantly increased when comparing the late latent infection (180 DPI) to lethal infection (**Figure 4A**). There was also a non-significant trend towards a decreased proportion of effector CD4 T-cells expressing the Th2 transcription factor gene *Gata3* (**Figure 4B**). No significant difference in the proportion of effector CD4 T-cells expressing the Th17 transcription factor gene *Rorc* or the Tfh transcription factor gene *Bcl6* was observed across all three experimental conditions (**Supplementary Figure 5**). However, there was a significant decrease in the proportion of effector CD4 T-cells expressing the Treg transcription factor gene *Foxp3* in the latent infections compared to the lethal infection (**Figure 4C**). Thus, these findings indicate an increase in Th1 polarization and decrease in Treg polarization over time during latent *C. neoformans* infection from 14 days post-infection to 180 days post-infection.

**Figure 4:**
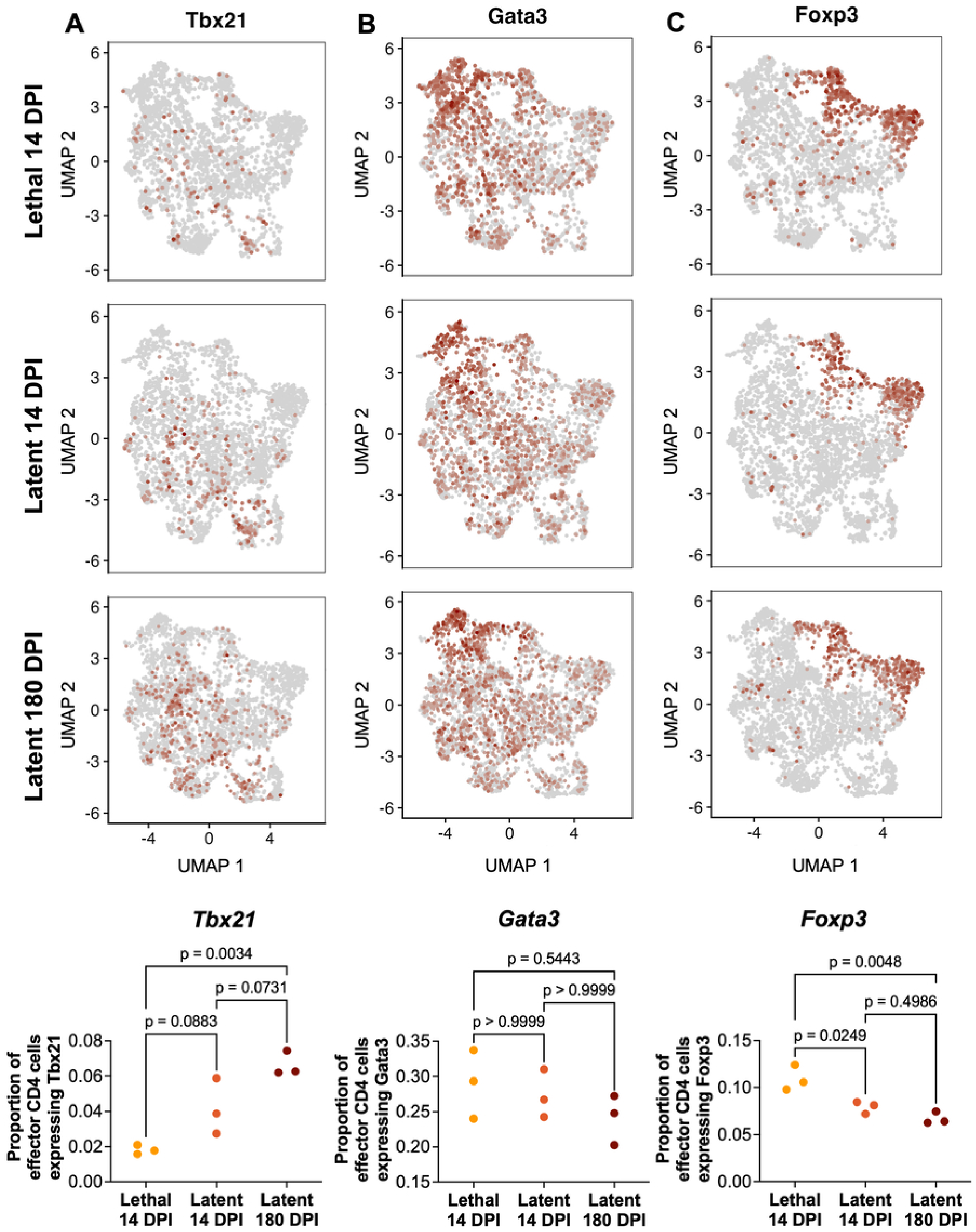
*Tbx21* (Tbet) expression by pulmonary effector CD4 T-cells is significantly increased during latent *C. neoformans* infection. Feature plots showing expression of (A) *Tbx21* (Tbet), (B) *Gata3* (GATA3), and (C) *Foxp3* (FoxP3) by pulmonary effector CD4 T-cells across the three experimental conditions: *C. neoformans* KN99*α* at 14 days post-infection (Lethal 14 DPI), UgCl223 at 14 days post-infection (Latent 14 DPI), and UgCl223 at 180 days post-infection (Latent 180 DPI). Cells expressing > 0 copies of the respective gene of interest were quantified and divided by the total number of effector pulmonary CD4 T-cells to determine the proportion of effector CD4 T-cells expressing the respective gene of interest in each biological replicate. Significance was determined via one-way ANOVA with Bonferroni correction.

Surprisingly, we also observed hybrid cells co-expressing multiple transcription factor genes (**Figure 5**). A higher proportion of cells co-expressing *Tbx21* (Tbet) and *Gata3* (GATA3) was observed in the latent infections compared to the lethal infection (**Figure 5A,D**). Cells co-expressing *Gata3* and *Rorc* (RORγt) and co-expressing *Tbx21* and *Rorc* were also identified in the effector subclusters (**Figure 5B-C**), but their prevalence did not differ across the various infections (**Figure 5D**).

**Figure 5:**
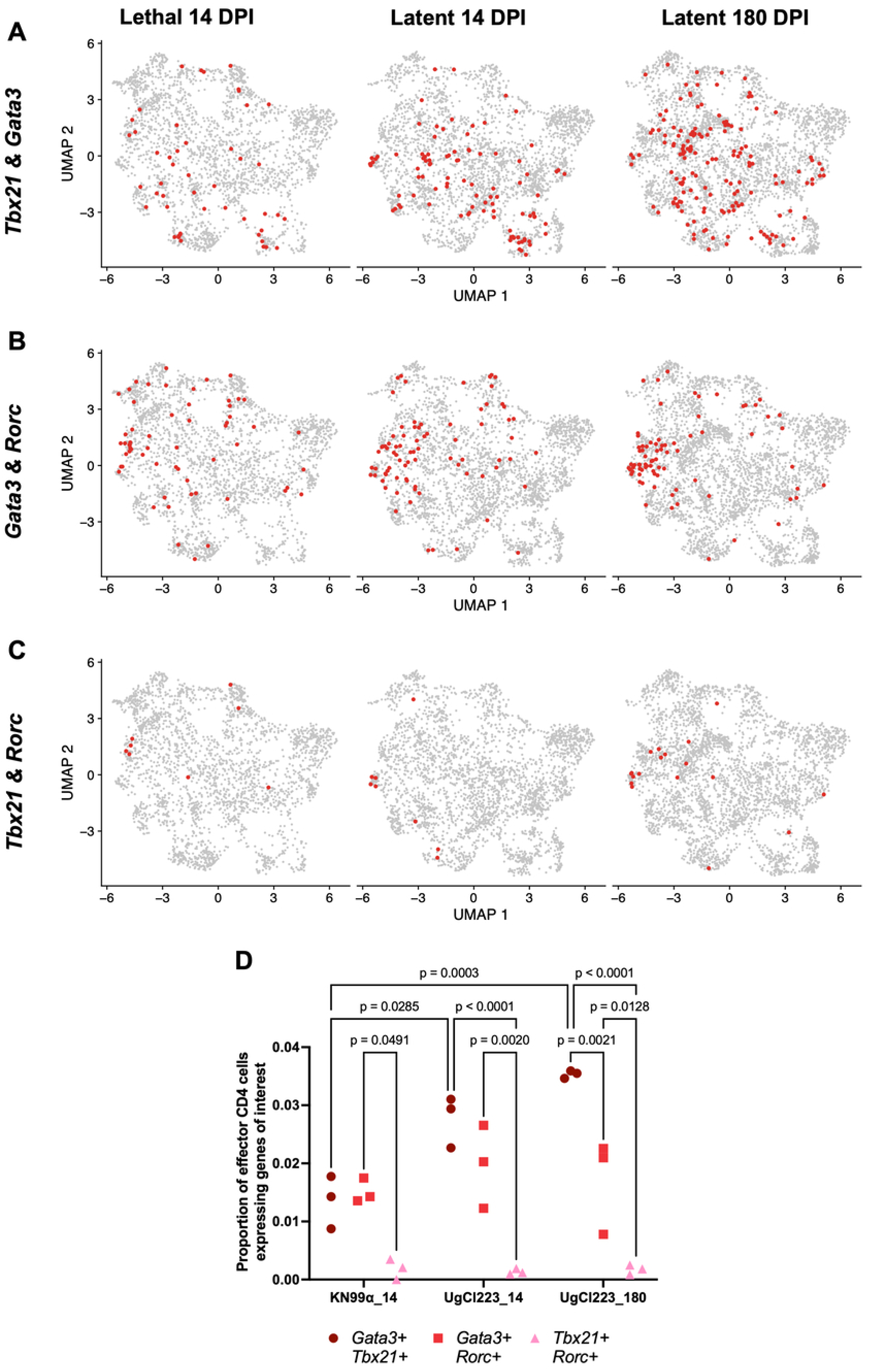
Co-expression of *Tbx21* (Tbet) and *Gata3* (GATA3) is significantly increased during latent *C. neoformans* infection. Feature plots showing co-expression of (A) *Tbx21* (Tbet) and *Gata3* (GATA3), (B) *Gata3* (GATA3) and *Rorc* (RORγt), and (C) *Tbx21* (Tbet) and *Rorc* (RORγt) by pulmonary effector CD4 T-cells across the three experimental conditions: *C. neoformans* KN99*α* at 14 days post-infection (Lethal 14 DPI), UgCl223 at 14 days post-infection (Latent 14 DPI), and UgCl223 at 180 days post-infection (Latent 180 DPI). (D) Cells expressing > 0 copies of the respective genes of interest were quantified and divided by the total number of effector CD4 T-cells to determine the proportion of effector CD4 T-cell co-expressing the respective genes of interest in each biological replicate. Significance was determined via two-way ANOVA with Bonferroni correction.

While Th1 polarization during latent infection increased from 14 to 180 DPI, there was still substantial heterogeneity at the 180 DPI timepoint. Cells expressing the Th1, Th2, Th17, and Tfh canonical transcription factors alone were identified (**Figure 6**; **Supplementary Table 1**). In addition, populations of hybrid cells expressing multiple transcription factors were also observed (**Figure 6**). Importantly, the largest population of cells in each infection were not identifiable based on expression of the canonical transcription factors (indicated as “other” in **Figure 6**). Preliminary analysis of this “other” population revealed the population was highly heterogeneous with expression of many genes that are not functionally characterized (data not shown). Taken together, these data show an upregulation of genes associated with Type-1 polarization in latent infections but also highlight a lack of known gene markers to characterize the entirety of the effector CD4 response to latent *C. neoformans* infection.

**Figure 6:**
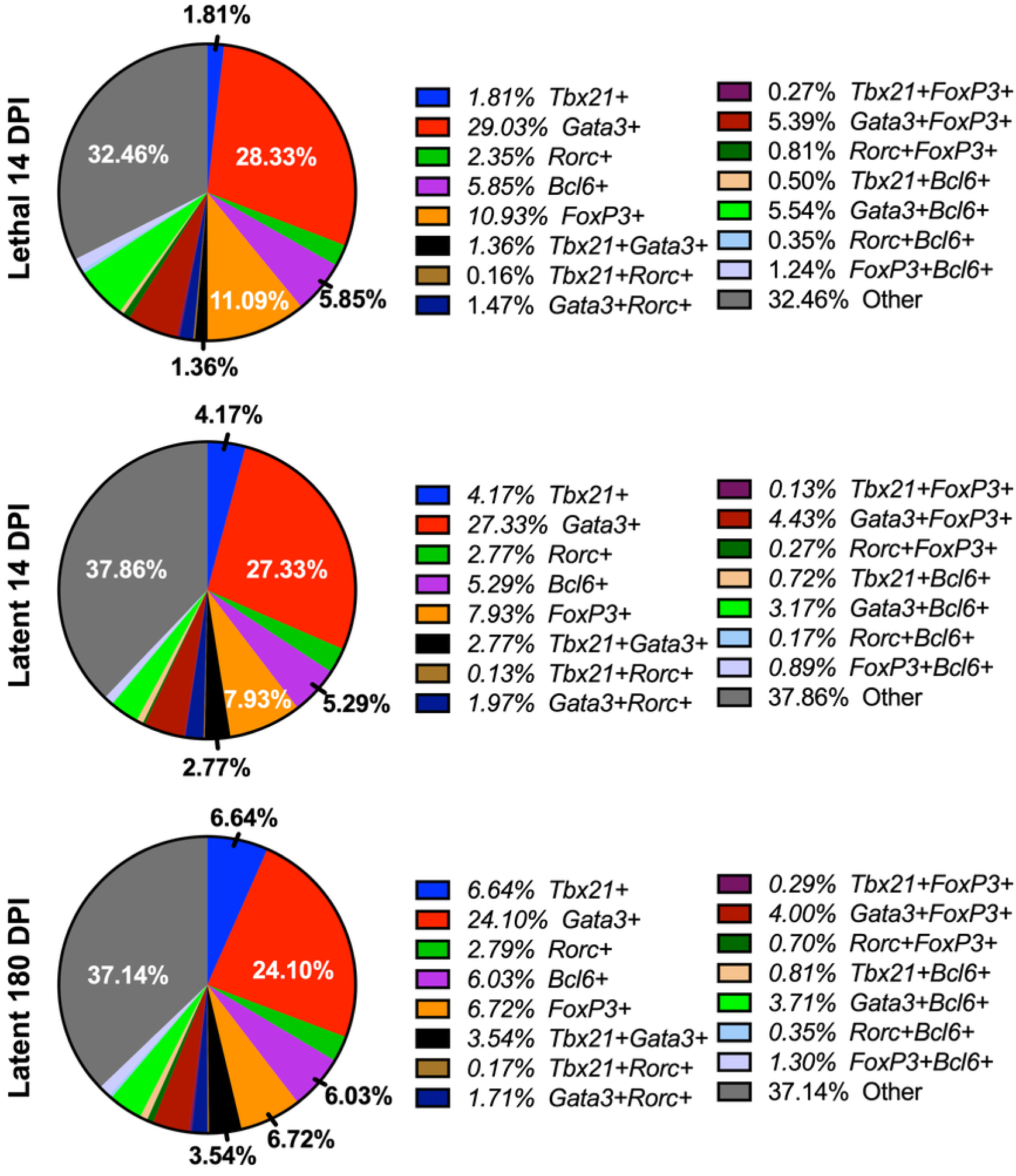
Pulmonary effector CD4 T-cell response to *C. neoformans* infection is highly heterogenous. Pie charts showing average percentage of pulmonary effector CD4 T-cells expressing lineage-defining transcription factor(s) across the three experimental conditions: *C. neoformans* KN99*α* at 14 days post-infection (Lethal 14 DPI), UgCl223 at 14 days post-infection (Latent 14 DPI), and UgCl223 at 180 days post-infection (Latent 180 DPI). Effector CD4 T-cells expressing > 0 copies of the transcription factor(s) of interest were identified via scRNAseq analysis, and percentage calculated from total number of effector CD4 T-cells. The remaining effector CD4 T-cells without known transcription factor expression were categorized as ‘Other’. Refer to **Supplementary Table 1** for standard error calculations.

### Adoptive transfer of Tbet+ CD4 T-cells can control latent infection in CD4-depleted mice

Given the enhanced Th1-like polarization observed during latent infection, we hypothesized that cells expressing the Th1 transcription factor *Tbet* are critical for controlling the latent infection. To isolate the effects of Tbet-expressing CD4 cells, lung resident Tbet-zsGreen-positive FoxP3-RFP-negative CD4 cells were isolated from the lungs of Tbet-zsGreen FoxP3-RFP mice latently infected with UgCl223 and then transferred into infection-matched CD4-depleted CD4-iDTR mice (henceforth referred to as ‘Tbet^pos^ adoptive transfer group’), and assessed for fungal burden (**Figure 7A**). Our results showed that the Tbet^pos^ adoptive transfer group had equivalent lung and brain fungal burden to the iDTR control mice (henceforth referred to as the ‘immune sufficient control group’) and had significantly lower lung and brain fungal burden when compared to the DT-treated CD4-iDTR mice latently infected with UgCl223 that did not receive any CD4 cells (henceforth referred to as ‘CD4-depleted control group’) (**Figure 7B-C**). Furthermore, 83% of the Tbet^pos^ adoptive transfer group survived until 49 days post-infection, compared to 75% of the CD4-depleted control group and 100% of the immune sufficient control group (**Figure 7D**). Overall, these data demonstrated that the Tbet-expressing CD4 T-cells were sufficient to control latent *C. neoformans* infection in mice.

**Figure 7:**
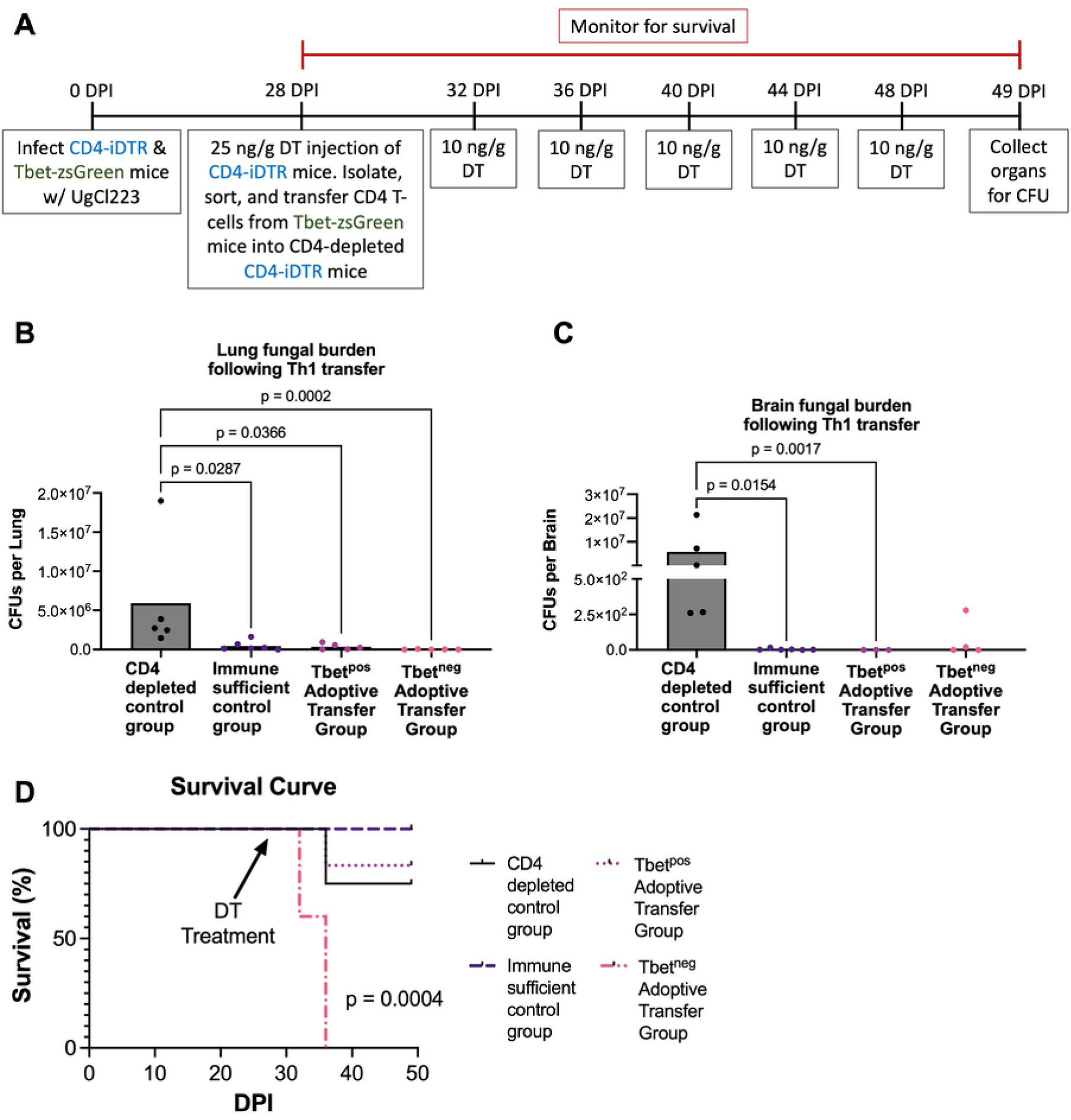
The pulmonary Th1-like response can control latent infection. (A) Tbet-zsGreen FoxP3-RFP mice and CD4-iDTR mice were latently infected with *C. neoformans* UgCl223. At 28 days post-infection (DPI), CD4-iDTR mice were injected with 25 ng/g diphtheria toxin (DT) via intraperitoneal (IP) injection. On the same day, lungs were isolated from infection-matched Tbet-zsGreen FoxP3-RFP mice, digested, negatively selected for CD4 T-cells, sorted for high zsGreen expression (Tbet^pos^ Adoptive Transfer Group) and low zsGreen expression (Tbet^neg^ Adoptive Transfer Group), and injected into the DT-treated CD4-iDTR mice. Every 4 days, the Tbet^pos^ Adoptive Transfer Group, Tbet^neg^ Adoptive Transfer Group, CD4-iDTR mice that were not injected with Tbet-zsGreen CD4 T-cells (CD4 depleted control group), and iDTR (immune sufficient control group) mice received a maintenance dose of 10 ng/g DT via IP injection. At 49 DPI, the control and Tbet^pos^ adoptive transfer groups were euthanized, and (B) lungs and (C) brain of mice were harvested, homogenized, and plated for colony forming units (CFUs). The Tbet^neg^ adoptive transfer group were euthanized and lungs and brain harvested at terminal endpoint (36 DPI). Significance was determined by one-tailed nonparametric t-test. (D) Mice were monitored for signs of morbidity and euthanized at terminal endpoint (20% total weight loss, 1 g/day weight loss for two consecutive days; or neurological symptoms including loss of sternal recumbency, partial paralysis, seizure, convulsion, or coma). Kaplan-Meier survival kinetics and significance was analyzed using log-rank testing.

To examine the effects of CD4 T-cells that did not express Tbet, we transferred lung resident Tbet-zsGreen-negative FoxP3-RFP-negative CD4 T-cells isolated from the lungs of Tbet-zsGreen FoxP3-RFP mice latently infected with UgCl223 into infection-matched CD4-depleted CD4-iDTR mice (henceforth referred to as ‘Tbet^neg^ adoptive transfer group’). Surprisingly, the Tbet^neg^ adoptive transfer group had 0% survival at 36 days post-infection (**Figure 7D**). The terminal lung and brain fungal burden was measured at this 36 DPI endpoint, prior to the 49 DPI timepoint used for the other mouse populations. Based on previous studies, we would anticipate no significant difference in fungal burden across the 36 and 49 DPI timepoints; and there was no difference in lung and brain fungal burden of the Tbet^neg^ adoptive transfer group compared to the Tbet^pos^ adoptive transfer group and the immune sufficient control group (**Figure 7B-C**). Instead, we observed a significant decrease in lung fungal burden in the Tbet^neg^ adoptive transfer group compared to the CD4 depleted control group (**Figure 7B**). These data suggest that, while the Tbet^neg^ cell population was able to control *C. neoformans* growth, these cells were unable to prevent mortality.

### IFNγ signaling is necessary for control of latent *C. neoformans* infection

Having demonstrated that Tbet+ cells were necessary and sufficient for controlling latent infection, we next aimed to clarify whether the downstream Th1-associated pathways, particularly IFNγ signaling, were similarly essential. *Ifngr1^+/+^* and *Ifngr1^-/-^* mice were latently infected with UgCl223 and monitored for survival (**Figure 8A**). We observed 100% mortality of the latently-infected *Ifngr1^-/-^* mice, with a median survival of 56 DPI (**Figure 8B**). These findings indicate that IFNγ signaling is a necessary component of the Th1 inflammatory response.

**Figure 8:**
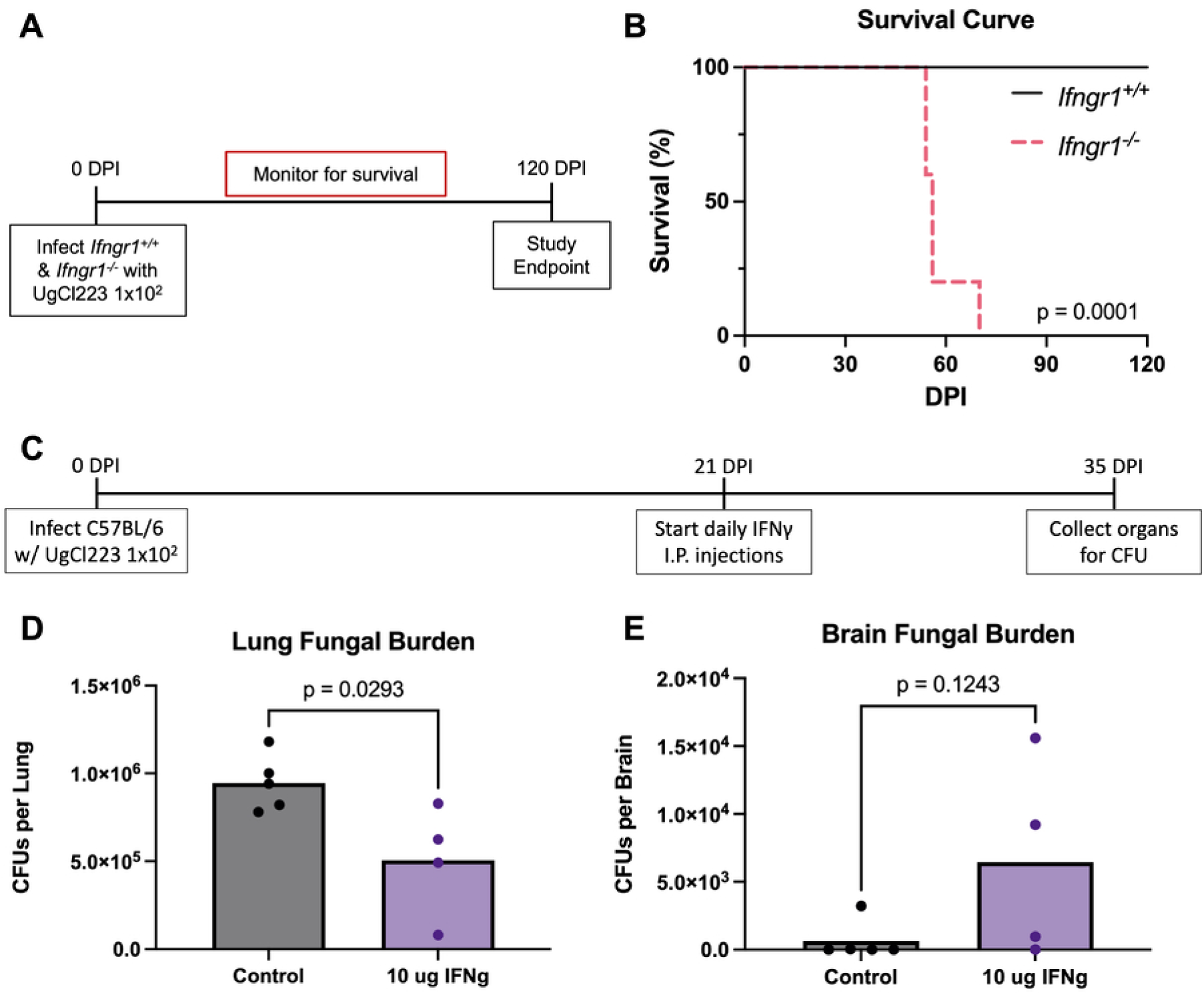
Exogenous IFNγ supplementation decreases lung fungal burden during latent *C. neoformans* infection. (A) *Ifngr1^+/+^* (i.e. C57BL/6J) and *Ifngr1^-/-^* mice were latently infected with *C. neoformans* UgCl223. (B) Mice were monitored for signs of morbidity and euthanized at terminal endpoint (20% total weight loss, 1 g/day weight loss for two consecutive days; or neurological symptoms including loss of sternal recumbency, partial paralysis, seizure, convulsion, or coma). Kaplan-Meier survival kinetics and significance was analyzed using log-rank testing. (C) C57BL/6J mice were infected with *C. neoformans* UgCl223. At 21 days post-infection (DPI), mice were injected with 10 *μ*g IFNγ or sterile PBS vehicle control via weekly intraperitoneal injections. At 35 DPI, (D) lungs and (E) brain of mice were harvested, homogenized, and plated for colony forming units (CFUs). Significance was determined by two-tailed t-test.

Interestingly, our previous study noted that pulmonary CD4 T-cells isolated from latently-infected mice had reduced IFNγ production when compared to cells isolated from uninfected control mice (12). We hypothesized that this diminished IFNγ production by the Tbet+ CD4 T-cells during latent infection prevents complete clearance of the infection and posited that exogenous IFNγ supplementation would enhance the antifungal response. To test this hypothesis, mice were latently infected with UgCl223 and injected daily with 10 μg of mouse recombinant IFNγ starting at 21 DPI. At 35 DPI (14 days post-IFNγ treatment), lung and brain fungal burden was assessed (**Figure 8C**). While the lung fungal burden in the IFNγ-treated mice was significantly decreased compared to that of sterile saline treated control mice, treatment with 10 μg was not sufficient to completely clear the infection from the lungs (**Figure 8D**). In addition, IFNγ treatment resulted in a non-significant trend towards increased brain fungal burden (p-value = 0.1243) (**Figure 8D**). Thus, IFNγ supplementation reduced lung fungal burden during latent infection, but did not completely resolve the infection.

### CTLA-4 expression was significantly increased during latent *C. neoformans* infection

Although we clearly established that Th1 polarization was beneficial in controlling pulmonary *C. neoformans* infection, we sought to reconcile this finding with the prominent inhibitory gene expression observed in our scRNAseq analysis of effector CD4 cells (cluster 2) (**Supplementary Figure 2**). Indeed, the proportion of effector CD4 cells expressing *Ctla4* was significantly increased compared to all other known regulators of immune exhaustion regardless of experimental condition (**Supplementary Figure 6**). The predominance of *Ctla4* expression was also reflected in our flow cytometry data, where the proportion of activated CD4+CD44+FoxP3- T-cells producing CTLA-4 was significantly increased throughout the latent infection, in comparison to cells isolated from uninfected mice (**Figure 9A**). In contrast, this phenomenon was not observed in activated CD4+CD44+FoxP3- T-cells producing PD-1, FoxP3, LAG-3, or TIM-3 (**Figure 9B-E**). Overall, the scRNA-seq and flow cytometry analysis both suggested that *Ctla4* (CTLA-4) could be integral to immunomodulation during latent infection, albeit with unclear implications.

**Figure 9:**
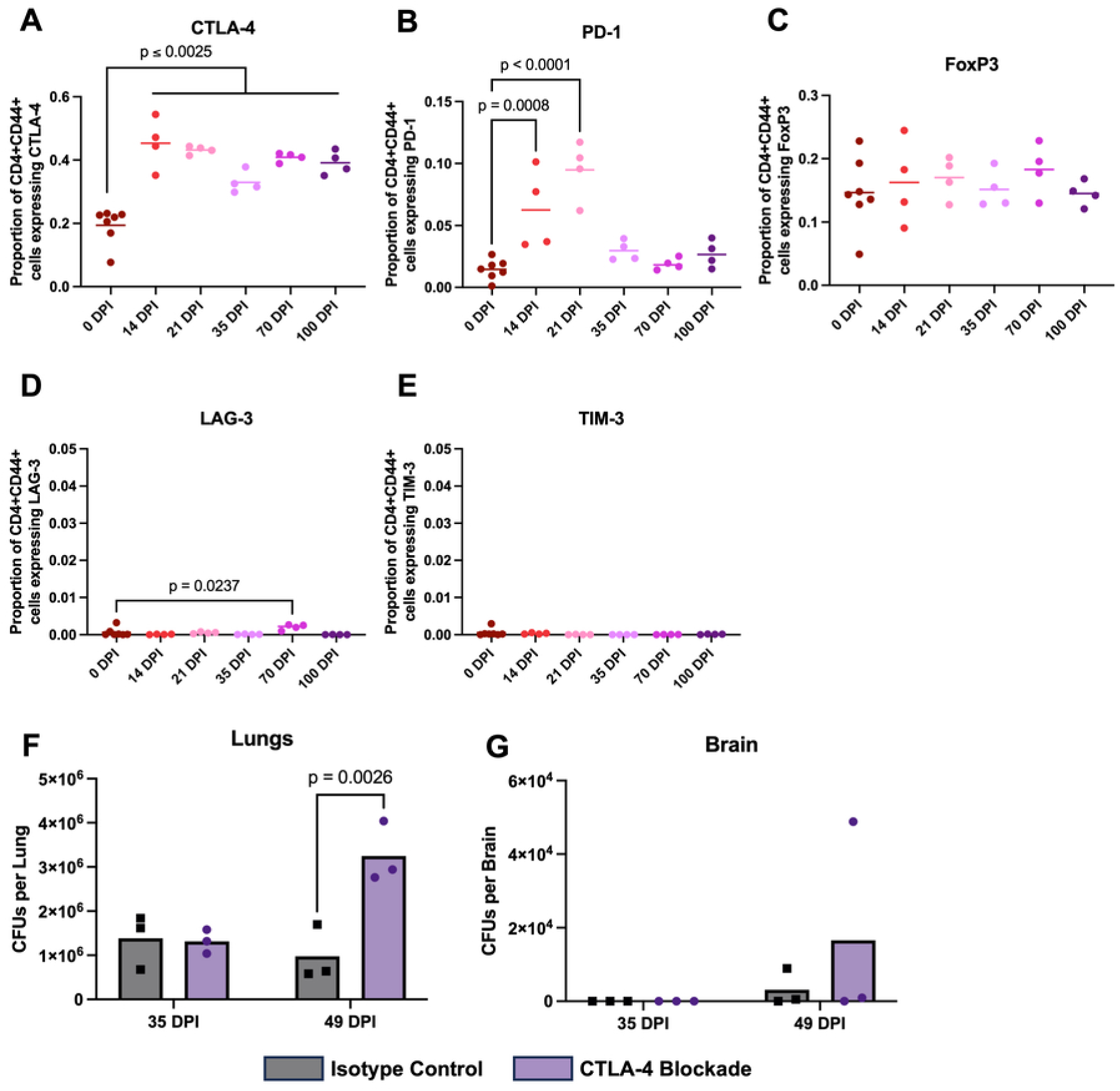
CTLA-4 blockade increases lung fungal burden. At 0-, 14-, 21-, 35-, and 100-days post-infection, lungs of C57BL/6J mice infected with *C. neoformans* UgCl223 were isolated and digested into single cell suspensions, and proportion of (A) CTLA-4+, (B) PD-1+, (C) LAG3+, (D) TIM3+, and (E) FoxP3+ expressing CD4+CD44+ cells were quantified via flow cytometry based on the total number of lung CD4+CD44+ cells. Significance was determined by one-way ANOVA with Bonferroni correction. For CTLA-4 blockade, C57BL/6J mice were infected with *C. neoformans* UgCl223 and at 21 days post-infection (DPI), mice were injected with 0.5 mg anti-CTLA4 monoclonal antibody (9D9) or IgG2b isotype control via weekly intraperitoneal injections. At 35- and 49- days post infection, (F) lungs and (G) brain of mice were harvested, homogenized, and plated for colony forming units (CFUs). Significance was determined by two-way ANOVA with Bonferroni correction.

Intriguingly within the scRNAseq data, there were differences in the expression of *Ctla4* amongst the effector CD4 cells that expressed either *Tbx21* (Th1), *Gata3* (Th2), or *Rorc* (Th17) across all experimental conditions (**Figure 10A-C**). For both lethal and latent infection at 14 DPI, the proportion of *Tbx21*-expressing effector CD4 T-cells that also expressed *Ctla4* was significantly lower than the proportion of *Gata3*-expressing and *Rorc*-expressing effector CD4 T-cells with *Ctla4* co-expression (**Figure 10D**). The only significant difference between the experimental conditions was an increase in the proportion of *Tbx21 Ctla4* co*-*expression between 14 DPI and 180 DPI (**Figure 9D**). These data indicate that *Ctla4* is more highly expressed by Th2 and Th17 cells, rather than Th1 cells, in early latent and lethal infection.

**Figure 10:**
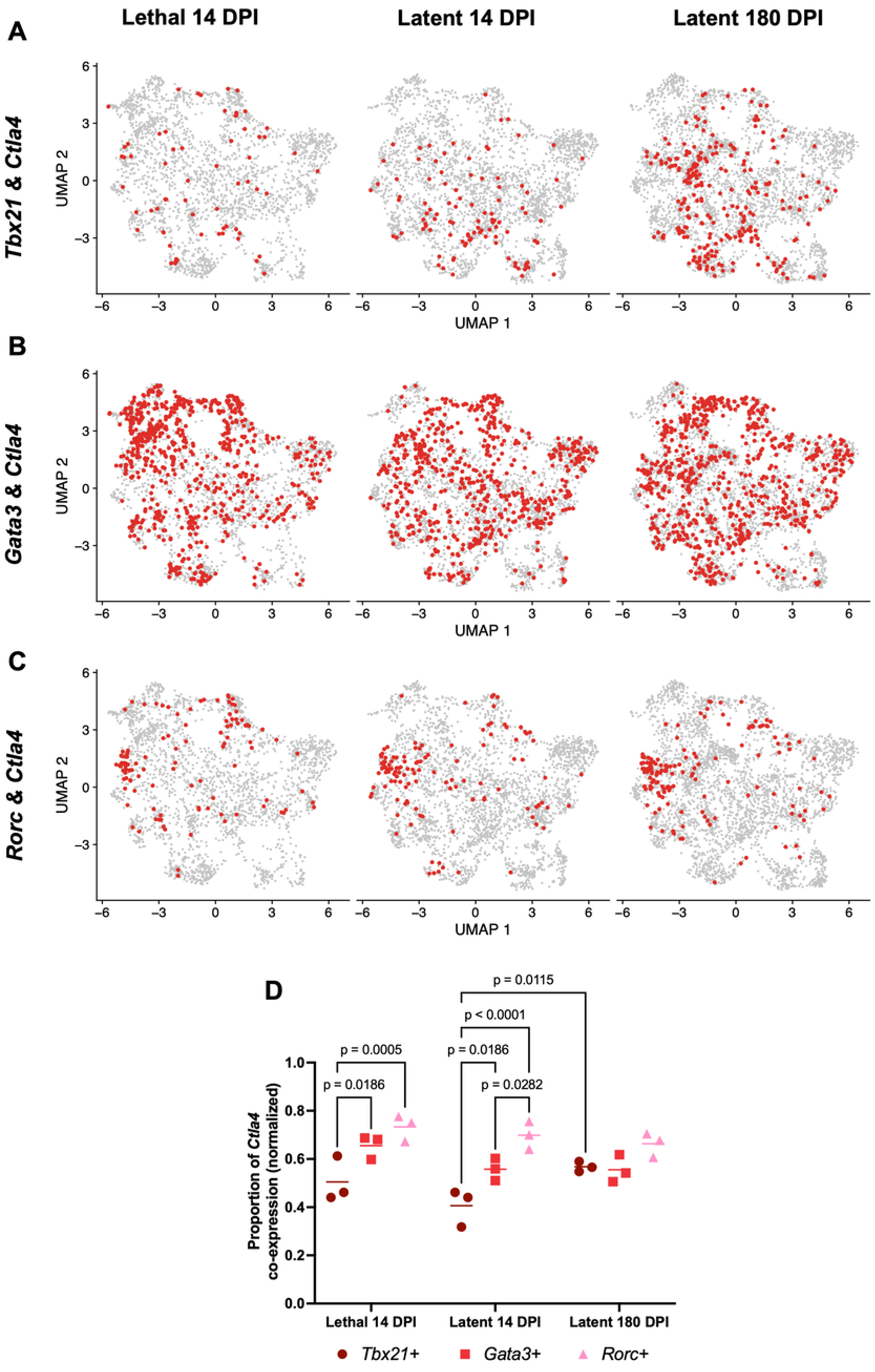
Co-expression of *Ctla4* (CTLA-4) and *Gata3* (GATA3) is significantly increased during *C. neoformans* infection. Feature plots showing co-expression of (A) *Tbx21* (Tbet) and *Ctla4* (CTLA-4), (B) *Gata3* (GATA3) and *Ctla4* (CTLA-4), and (C) *Rorc* (RORγt) and *Ctla4* (CTLA-4) by pulmonary effector CD4 T-cells across the three experimental conditions: *C. neoformans* KN99*α* at 14 days post-infection (Lethal 14 DPI), UgCl223 at 14 days post-infection (Latent 14 DPI), and UgCl223 at 180 days post-infection (Latent 180 DPI). (D) Effector CD4 T-cells co-expressing > 0 copies of *Ctla4* and > 0 copies of *Tbx21*, *Gata3*, or *Rorc* were quantified and divided by the total number of cells expressing *Tbx21*, *Gata3*, or *Rorc*, respectively, to determine the proportion of each cell population with *Ctla4* co-expression in each biological replicate. Significance was determined via two-way ANOVA with Bonferroni correction.

To determine the role of CTLA-4 during latent infection, we tested whether blockade of CTLA-4 influenced fungal proliferation in the latent infection model. Following latent infection with UgCl223, mice were injected weekly with anti-CTLA-4 monoclonal antibody starting at 21 DPI. At 35 and 49 DPI, lung and brain fungal burden were assessed. CTLA-4 blockade promoted fungal proliferation, with significantly higher lung fungal burden in mice treated with the CTLA-4 monoclonal antibody compared to mice treated with an isotype control at 49 DPI (**Figure 9F**). We also observed a non-significant increase in brain fungal burden between the CTLA-4 blockade and the isotype control mice (**Figure 9G**). Altogether, these data support a protective role for CTLA-4 in the context of early latent *C. neoformans* infection.

## DISCUSSION

CD4 T-cells are a necessary host defense against *C. neoformans* infection, as evidenced by the significant global burden on HIV-associated cryptococcal meningitis (1). These observations are also well-corroborated by *in vivo* animal depletion studies that used monoclonal antibodies (52, 53) and diphtheria toxin-induced cell lineage ablation (12) which showed that CD4 T-cells are necessary for pulmonary control of *C. neoformans* infection. In this study, we further characterized the effector CD4 T-cell response via scRNAseq analysis and identified key mechanisms of host control against latent *C. neoformans* infection.

Using scRNAseq, we explored the transcriptional landscape of the CD4 T-cell response against both lethal and latent *C. neoformans* infections. We identified 10 distinct CD4 T-cell subsets, of which 9 were naïve CD4 T-cells and the remaining cluster was annotated as effector CD4 T-cells (cluster 2) based on upregulation of *Cd44*. Notably, the top 10 expressed genes for each naïve CD4 T-cell cluster were unique to each individual cluster, which highlighted the heterogeneity of the pulmonary naïve CD4 response. The expression of genes involved in T-cell activation such as *Camk1d* and *Ifit3*/*Ifit1* also suggested that the naïve CD4 T-cell repertoire was influenced by the *C. neoformans* infection (15, 16).

Following the identification of the effector CD4 cluster, we performed additional clustering analysis and revealed 12 effector CD4 subclusters. As with the naïve CD4 T-cell clusters, annotation of the effector CD4 subclusters was limited to well-characterized genes. However, by using pseudotime analysis, we were able to further infer the trajectories of CD4 T-cell activation and differentiation from the “newly arrived” lymph node-derived CD4 T-cells. This pseudotime analysis, when combined with the differential gene expression analysis of the effector CD4 subclusters, revealed a significant increase in Th1 polarization, and decrease in Th2 and Th17 polarization, over the time course of the latent infection. These findings are consistent with our previous studies that showed the immune response during latent infection changes from a preliminary Th2, Th17 response to the mature Th1 response (51). Thus, we now have strong evidence that links latent *C. neoformans* infection with increased Th1 polarization.

Importantly, the scRNAseq analysis of pulmonary CD4 T-cells during *C. neoformans* infection highlighted the immense transcriptional heterogeneity of the CD4 T-cell response and revealed that a substantial proportion (∼30%) of effector CD4 T-cells could not be identified based on known transcription factor production. These cells express various poorly characterized genes, thus it remains unclear what role these undefined CD4 T-cells play during latent *C. neoformans* infection. The identification of hybrid cell populations is also intriguing and suggests more nuanced regulation of the phenotype of Th cells that previously appreciated. Importantly, the diversity among the CD4 T-cells we observed is predominantly within the “Lung CD4+ TD” population in the newly proposed T-cell nomenclature and highlights the challenges related to classifying the diversity of CD4+ T-cell phenotypes during infection and identification of key cell populations (54, 55).

By comparing 1) lethal and latent infections at the same timepoint and 2) early and late latent infection timepoints we gained a better understanding of the Th cell phenotypes that influence disease outcomes in *C. neoformans*. Lethal infections promoted a strong Th2 response with significant upregulation of immunosuppressive regulatory genes in the majority of effector subclusters. These data are consistent with previous studies showing Th2 responses dominate during lethal *C. neoformans* infections (8, 9). Importantly, the significant increase in proportion of effector CD4 T-cells expressing *Tbx21*, or co-expressing *Tbx21* and *Gata3*, during latent infection corroborates the damage-response framework theory (56–58), where the host response to *C. neoformans* infection oscillates between a beneficial controlled Type-1 polarization versus a detrimental Type-2 polarization.

Surprisingly, the Tbet+ Th1-like CD4 T-cells only account for 10% of the total effector cell population. Yet the adoptive transfer of pulmonary Tbet+ CD4 cells proved that Tbet+ CD4 T-cells are necessary and sufficient to control latent *C. neoformans* infection. Of note, this experimental set-up was not able to differentiate between CD4 T-cells that only express Tbet+ and hybrid cells that co-express Tbet+ and other lineage defining transcription factors. Regardless, the significant decrease in fungal burden and improved survival of the Tbet^pos^ adoptive transfer group compared to the CD4-depleted group suggests that the diverse Tbet+ CD4 populations were both necessary and sufficient for preventing proliferation and dissemination, and ultimately mortality during *C. neoformans* latent infection.

With regard to the Tbet^neg^ adoptive transfer group, the low fungal burden observed during this experiment suggests that the Tbet^neg^ CD4 cells were at least partially capable of controlling the fungal infection. We cannot rule out the possibility that this decreased fungal burden simply reflects the earlier timepoint (36 DPI) of organ collection compared to the other experimental and control groups (49 DPI), however previous latent infections studies show no significant changes in fungal burden between 36 and 49 DPI (12, 51). However, the increased mortality of mice receiving the Tbet^neg^ adoptive transfer is striking and suggests these cells could act as a double-edged sword; helping control fungal proliferation yet eliciting rapid mortality. Of note, the design of the adoptive transfer experiments excluded Tregs, which may be critical for dampening or controlling the activity of the diverse Tbet^neg^ effector CD4 T-cell subsets. It is also conceivable that the beneficial activities of the undefined CD4 T-cell subsets were masked by the function of Gata3+ Th2-like CD4 T-cells, which were previously shown to induce detrimental Type-2 responses during lethal infections (9).

In addition to demonstrating that Tbet+ cells are necessary and sufficient for controlling latent infection, we also showed that downstream IFNγ signaling is a key component, with 100% of latently-infected *Ifngr1^-/-^* mice succumbing to the infection. Furthermore, we observed a decrease in lung fungal burden with exogenous IFNγ treatment in latently-infected mice. However, IFNγ supplementation did not completely clear latent infection, in contrast to previous studies where infection with a *C. neoformans* strain genetically modified to produce IFNγ resolved without additional interventions (59, 60). It is certainly possible that the diminished IFNγ response observed during latent infection (12, 61) may reflect *C. neoformans*-mediated modulation of the host response and long-term persistence within the lungs; but our findings reflect the difficulties in determining an appropriate therapeutic dose for IFNγ. These conclusions are not surprising, as adjunctive IFNγ therapy did not impact survival outcome between treated and untreated patient cohorts, although a decrease in brain fungal burden was observed in these human clinical trials (62). While a higher dose of IFNγ may theoretically result in greater fungal clearance, the trend towards increased brain fungal burden with IFNγ supplementation that we observed suggests that IFNγ may enhance either dissemination or growth of *C. neoformans* in the brain via a phenomenon similar to immune reconstitution inflammatory syndrome (IRIS) (63). Thus, CD4 T-cell production of IFNγ is important for controlling *C. neoformans* infection, but its utility as a therapeutic agent may be limited.

To date, the scientific community has focused on understanding the effects of Th1 polarization on *C. neoformans* infection. However, our scRNAseq analysis of the effector CD4 T-cell response to *C. neoformans* infection revealed upregulation of many immune regulatory genes, most significant of which was *Ctla4* (CTLA-4) (64, 65). Indeed, significant upregulation of CTLA-4 was observed via flow cytometry as early as 14 DPI during latent *C. neoformans* infection, which reflects previously published data demonstrating that the capsule of *C. neoformans* strains can promote rapid expression of CTLA-4 in CD4 T-cells within 24 hours of co-incubation (66).

Interestingly, we found that the proportion of *Gata3*-expressing Th2-like cells co-expressing *Ctla4* was significantly higher than that of the *Tbx21*-expressing Th1-like cells. Furthermore, CTLA-4 monoclonal antibody (mAb) blockade resulted in significantly increased lung fungal burden. These findings are in contrast to a previous study that demonstrated CTLA-4 blockade promotes fungal clearance during lethal *C. neoformans* infection (67). The opposing experimental outcomes may be due to multiple experimental factors, including the type of *C. neoformans* strain, timing of the administration of the CTLA-4 mAb, and different inbred mouse strains. In addition, differences in host CD4 polarization against lethal and latent *C. neoformans* infection likely also contributed to differences in fungal clearance or proliferation. Regardless, our data suggest that CTLA-4 mAb blockade starting at 21 DPI enhanced fungal proliferation and lead us to hypothesize that the blockade removed CTLA-4 suppression of the detrimental Th2 effector function.

There are still some points of uncertainty regarding CTLA-4 function during *C. neoformans* infections. For instance, we observed a significant increase in the proportion of *Ctla4* expression in Th1 cells (expressing *Tbx21*) from 14 to 180 DPI during latent infection, but the implication of this finding is unclear given that there was no difference in Ctla4 expression across the cell types at 180 DPI. In addition, while Th17 cells make up a small proportion of effector CD4 T-cells during *C. neoformans* infection, the proportion of *Ctla4* expression in Th17 cells (expressing *Rorc*) was significantly higher compared to that of Th1 and Th2 cells during lethal and early latent infections. These data suggest that studies to better delineate the role of Th17 cells are warranted, especially during latent infection.

In summary, we demonstrated that the CD4 T-cell response generated against *C. neoformans* infection is highly complex and heterogenous. We definitively showed that Tbet+ cells are necessary and sufficient to control latent *C. neoformans* infection. Our findings also shed light on the nuances that drive the host response to latent versus lethal *C. neoformans* infection, with special consideration to the integral role of immune regulation. We propose that the beneficial host response that leads to control during latent *C. neoformans* infections is paradoxical – involving both accumulation of beneficial Tbet+ cells and a simultaneous CTLA-4 mediated suppression to dampen activity of other cell types. Our data and previous studies suggest targeting IFNγ may not be a viable clinical option. However, CTLA-4 activation has not been explored and could be a novel immunomodulatory mechanism to prevent cryptococcosis.

## MATERIALS & METHODS

### Ethics statement

Animal experiments were done in accordance with the Animal Welfare Act, United States federal law, and National Institutes of Health guidelines. Mice were handled in accordance with guidelines defined by the University of Minnesota Animal Care and Use Committee (IACUC) under protocols 1908A37344, 2207A40205, and 2104A39016; and the National Institutes of Health IACUC under protocol LHIM4E.

### Inbred mouse strains

All mice used in this study were C57BL/6J, or derived from a C57BL/6J background. Mice were housed in specific pathogen-free conditions. CD4-Cre (68) and homozygous iDTR (69) breeding pairs were purchased from Jackson Laboratory and were crossed to generate CD4-Cre/iDTR, referred to as CD4-iDTR, mice (69). Systemic CD4 T-cell ablation in the CD4-iDTR mice was achieved by an initial intraperitoneal injection of 25 ng/g diphtheria toxin (DT; Sigma) followed by an additional 10 ng/g DT every 4 days. CD4+ T cell levels were monitored to confirm the effectiveness of the DT treatment throughout the 49 DPI treatment regimen. Tbet-zsGreen mice (70) were crossed with FoxP3-RFP mice (71) to generate Tbet-zsGreen FoxP3-RFP mice and were kindly gifted by Marc Jenkins. *Ifngr1^-/-^* mice were obtained from Jackson Laboratory (JAX 025545) (72). Mice used for all experiments were 6-8 weeks of age; all controls were sex- and age-matched.

### Cryptococcus strains

*Cryptococcus neoformans* strains KN99α (73) and UgCl223 (74, 75) were stored as -80°C glycerol stocks, streaked on yeast peptone dextrose (YPD) + 0.04 g/L chloramphenicol agar plates and incubated for 2 days at 30°C prior to use. YPD broth was inoculated with colonies from the aforementioned plate and incubated for 16 hours at 30°C and 225 RPM. The resulting inoculum was prepared by centrifuging the culture for 1 minute at 14,000 RPM (17,968 x g) to pellet the cells, washing the cells 3 times with phosphate buffered saline (PBS), and resuspending the cells in PBS at a concentration of either 1x10^6^ cells/mL (KN99α) for lethal infections or 2x10^3^ cells/mL (UgCl223) for latent infections.

### Infection

A well-established intranasal pulmonary inhalation model of cryptococcosis was used for this study (73, 76). Mice were fully anesthetized with pentobarbital until they did not respond to a toe pinch. The mice were then lethally infected with 5x10^4^ KN99α or latently infected with 1x10^2^ UgCl223 *C. neoformans* cells in 50µL of PBS by placing the inoculum on the nares of each mouse. The mice were then suspended by their incisors for 5 minutes and subsequently paced upright on a paper towel in their cage until regaining consciousness, enabling the inoculum to reach the lower respiratory tract.

All mice were monitored for morbidity and sacrificed at pre-determined timepoints or when endpoint criteria were reached. Endpoint criteria were defined as 20% total body weight loss, loss of 2 grams of weight in 2 days, or symptoms of neurological disease.

### Organ fungal burden

Lungs and brain were collected at the time of mouse euthanasia and placed into 2 mL sterile PBS. The collected tissues were homogenized, serial dilutions of tissue homogenates were plated on YPD agar plates supplemented with 0.04 mg/mL chloramphenicol and incubated at 30°C. *C. neoformans* colonies were counted after 48 hours of incubation.

### Single cell RNA sequence workflow and analysis

We conducted two separate single cell RNA sequencing experiments. 1) A preliminary study analyzing total lung CD45+ leukocytes isolated from a mouse infected with *C. neoformans* UgCl223 at 14 days post-infection, a mouse infected with UgCl223 at 150 days post-infection, a mouse infected with *C. neoformans* KN99α at 14 days post-infection, and an uninfected mouse. 2) A robust study analyzing lung CD4+ lymphocytes isolated from three mice infected with UgCl223 at 14 days post-infection, three mice infected with UgCl223 at 180 days post-infection, and three mice infected with KN99α at 14 days post-infection.

For both studies, lungs were perfused with cold PBS and digested into a single cell homogenate with Collagenase A. For the CD45+ preliminary study, CD45+ cells were isolated via positive selection using the EasySep™ Mouse CD45 Positive Selection Kit (StemCell) per manufacturer’s instructions. For the CD4+ study, lymphocytes were first isolated via a 67%/40% Percoll density gradient, and then CD4 T-cells were isolated via negative selection using the EasySep™ Mouse CD4+ T cell Isolation Kit (StemCell) (see Flow cytometry methods subsection below for a more detailed description).

For both studies, single cell RNA sequencing libraries were prepared using a Chromium Single Cell Controller (10x Genomics). The CD45+ preliminary study used the Chromium Single Cell 3’ Library and Gel Bead v3.1 dual index kit (10x Genomics). The CD4+ study used the Chromium Single Cell 5’ Library and Gel Bead v2 dual index kit (10x Genomics). Briefly for both studies, cell suspensions were diluted in nuclease-free water to obtain a target cell recovery of 10,000 cells, according to manufacturer’s instructions. Remaining steps were also carried out according to the manufacturer’s instructions for cDNA amplification and sample index PCR. For both studies, final libraries were sequenced on two NovaSeq 6000 (Illumina) 150-bp paired end flow cells at the University of Illinois at Urbana-Champaign DNA Services Lab. Libraries were sequenced with read lengths of 28x10x10x150. For both studies, the *Mus musculus* assembly mm10-2020-A (GRCm38) from NCBI were used with Cell Ranger 7.0.1 to make a reference for alignment. The ‘cellranger count’ command was then used to call cells and count UMIs for each gene and cell.

For both studies, UMI count matrices were imported into R from the Cell Ranger output. The CD45+ preliminary study was performed with R (version 4.0.4) and Seurat (version 4.3.0.1). The CD4+ study was performed with R (version 4.4.0) and Seurat (version 5.0.3). Genes were discarded if there were fewer than 10 cells with UMIs (unique reads). Counts were then imported into a Seurat object. Percent mitochondrial UMIs (“percent.mt”) were calculated using the thirteen mitochondrial genes in the Cell Ranger reference. The percentage mitochondrial UMIs, along with total UMIs per cell and total features per cell with UMIs were recorded and are available upon request. Cells with fewer than 200 features with UMIs were removed, as they were assumed to be red blood cells, low quality, or dying cells. Cells with more than 3 median absolute deviations above the median number of UMIs were filtered, given that they were likely to represent doublets. Cells with more than 3 median absolute deviations above the median number of mitochondrial UMIs were discarded, because they were assumed to be dying. UMI counts were normalized via the SCTransform method (77) using the top 3,000 most variable features and regressing on percentage mitochondrial UMIs and cell cycle scores.

For the CD45+ preliminary study, all samples were integrated using an anchor-based canonical correlation analysis (CCA) approach (78). Principal component analysis (PCA) was performed on the transformed UMI counts, and the first 40 axes were selected by the heuristic elbow method, and were retained for clustering and Uniform Manifold Approximation and Projection (UMAP). Following UMAP visualization, calculation of k-nearest neighbors was performed. Clustering analysis was used to determine the appropriate resolution and number of clusters for each experimental condition (79–81). Seurat was used for identifying cell clusters (82). Microarray data from Immunological Genome Project (83), distributed with the celldex R package (84), was used. Low quality assignments were pruned according to default settings. Following the annotation of each Seurat cluster, differential expression analysis was performed by grouping the infected experimental conditions and comparing against the uninfected condition via a Model-based Analysis of Single-cell Transcriptomics (MAST) approach (zero-inflated negative binomial model) (85). Genes with adjusted p-values <0.05 for each comparison were ranked by log2FC.

For the CD4+ study, we merged all experimental groups together, but opted against performing integration across the different replicates and experimental groups since we did not observe condition-specific clustering in our merged dataset (**Figure 1D**). Normalization, PCA, UMAP, and clustering analysis were performed as described above. To increase the rigor of our study, we identified clusters with elevated *Cd4* expression and excluded the clusters with low expression of *Cd4* and cell clusters with non T-cell markers (i.e. endothelial, alveolar, fibroblasts). In addition, to reduce artificial cluster separation caused by *Trbv* gene overrepresentation, all *Trbv* genes were removed from the variable gene set and all subsequent sub-clusterings. Normalization, dimensionality reduction, and clustering analysis were repeated to identify all *Cd4* positive cell clusters. Cluster-specific markers were identified using FindAllMarkers().

Clusters were annotated based on the top 10 expressed genes. Of the 10 Cd4 clusters, only cluster 2 had high expression of *Cd44* (**Figure 1B**), identifying it as the Effector Cd4 cluster. Cluster 2 (Effector) was isolated from the larger *Cd4* positive dataset for subclustering analysis, which included further normalization, dimensionality reduction, and clustering analysis. We opted to perform integration analysis during the subclustering analysis for better alignment of the Cluster 2 (Effector) subclusters. Differential expression analysis of the Cluster 2 (Effector) subclusters was performed using muscat and limma. A model matrix was constructed to compare conditions pairwise (Lethal 14 DPI versus Latent 14 DPI; Latent 180 DPI versus Latent 14 DPI). Trajectory analysis was conducted using Monocle3 (v1.3.1). Cells were ordered in pseudotime, with trajectory root manually inferred from Subcluster 1.

All sequencing data were deposited at NCBI under accession number PRJNA1271018 and the bioinformatic analysis was deposited in Dryad at https://doi.org/10.5061/dryad.j3tx95xsx.

### Flow cytometry

We utilized intravascular staining to discriminate between vascular and tissue leukocytes as described previously (86). Briefly, mice were intravenously injected via tail vein with APC-eFluor780-labelled CD45 antibody (30-F11, eBioscience) and were euthanized three minutes following injection with CO_2_. Pulmonary leukocytes were isolated as described previously (87). Briefly, the chest cavity was opened, cold PBS was used to perfuse lungs via right ventricle, and the trachea was exposed. Lungs were inflated with 2 mL of digestion solution containing 1.5 mg/mL Collagenase A (Roche), 5 mM DNase I (Ambion), 5% fetal bovine serum (FBS), and 10 mM HEPES (MP Biomedicals) in Hanks’ Balanced Salt Solution (HBSS, Gibco). Lungs were excised and placed in 5 mL digestion solution. Lung tissue plus digestion solution was incubated in a 37°C water bath for 30 minutes with gentle vortexing every 8-10 minutes. Upon completion of digestion, the resulting cell suspensions were strained through a 70 μM cell strainer and washed with 25 mL of PBS. Cells were then centrifuged for 10 minutes at 300 x g for 10 minutes at 4°C.

For CD4 T-cell isolation, the resulting cell pellet was resuspended in 40% Percoll-RPMI medium (GE Life Sciences). A Percoll density gradient was created (5mL of 40% Percoll-RPMI top, 3mL 67% Percoll-PBS bottom) and the samples were centrifuged for 20 minutes at 650 x g without braking. The lymphocytes at the interface were removed, washed twice with PBS containing 1 mg/mL bovine serum albumin (BSA, Sigma) and 0.002 mM EDTA (Invitrogen), and then centrifuged for 10 minutes at 300 x g for 10 minutes at 4°C. CD4 T-cells were isolated via negative selection using the EasySep™ Mouse CD4+ T cell Isolation Kit (StemCell).

All single-cell suspensions were stained with Near-IR Live Dead viability dye (Biolegend) according to manufacturer’s instructions and then incubated for 15 minutes on ice with CD16/32 antibody (Biolegend) to prevent nonspecific antibody binding. For surface staining, samples were stained with fluorophore-labelled antibodies at 4°C for 30 minutes.

For intracellular staining, cells were washed, fixed, and permeabilized using the Foxp3/Transcription Factor Staining Buffer Set (eBioscience) according to the manufacturer’s instructions. Samples were stained intracellularly with fluorophore-labelled antibodies at 4°C overnight. After staining, cells were washed with 1x permeabilization buffer, placed in cell staining buffer, and stored at 4°C until ready for data acquisition by flow cytometry.

All data was acquired with a BD LSR Fortessa flow cytometer using BD FACSDiva software (BD Bioscience). Compensation was performed at the beginning of each experiment with UltraComp eBeads plus Compensation Beads (Invitrogen) and the ArC Amine Reactive Compensation Bead Kit (Life Technologies). Data was analyzed using FlowJo v10.8.1.

For CD4 exhaustion maker identification, the following fluorophore-labelled antibodies were used: B220 (RA3-6B2, APC-eFluor780, Thermo Fisher Scientific), CD11c (N418, APC-eFluor780, Thermo Fisher Scientific), CD11b (M1/70, APC-eFluor780, Thermo Fisher Scientific), F4/80 (BM8, APC- eFluor780, Thermo Fisher Scientific), CD3 (17A2, BV650, Biolegend), CD4 (GK1.5, BUV39, BD Biosciences), CD8 (53-6.7, AF700, Thermo Fisher Scientific), CD44 (IM7, PerCp-Cy5.5, Biolegend), PD-1 (J43, APC, Invitrogen), LAG-3 (C9B7W, BV421, Biolegend), TIM-3 (RMT3-23, BV711, Biolegend), CTLA-4 (UC10-4B9, BV605, Biolegend), and FoxP3 (FJK-16S, AF488, eBioscience). The gating strategy is shown in **Supplementary Figure 7**.

### Adoptive Transfer

Tbet-zsGreen FoxP3-RFP mice and CD4-iDTR mice were infected with *C. neoformans* UgCl223 as described above. At 28 days post-infection, CD4-iDTR mice were injected with 25 ng/g diphtheria toxin via intraperitoneal injection. On the same day, Tbet-zsGreen FoxP3-RFP mice were injected with CD45-APC via tail vein to differentiate between organ-specific and vascular cells and euthanized after 3 minutes. Lungs were perfused with cold PBS and digested via collagenase A into a single cell suspension (as described above) and stained for CD3 (17A2, AF700, Biolegend). Lung cells from 10 Tbet-zsGreen FoxP3-RFP mice were pooled together. Following negative CD4 isolation, cells were sorted for viability and the population of cells that were 1) CD45-APC- (lung resident), CD3-AF700+, FoxP3-RFP-, and Tbet-zsGreen+ and 2) CD45-APC- (lung resident), CD3-AF700+, FoxP3-RFP-, and Tbet-zsGreen- were isolated. The sorted lung resident Tbet-zsGreen+ and lung resident Tbet-zsGreen- cells were then injected into DT-treated CD4-iDTR mice (Tbet^pos^ adoptive transfer group and Tbet^neg^ adoptive transfer group, respectively). Every 4 days, the Tbet^pos^ adoptive transfer group (n = 6), the Tbet^neg^ adoptive transfer group (n = 5), CD4-iDTR mice that were not injected with Tbet-zsGreen CD4 T-cells (CD4 depleted control group) (n = 8), and iDTR mice (Immune sufficient control group) (n = 6) received a maintenance dose of 10 ng/g DT via IP injection. At 49 DPI, surviving experimental and control groups were euthanized. Lungs and brains were assessed for fungal burden as described above.

### Ifngr1^-/-^ survival curve and exogenous IFNγ supplementation

*Ifngr1^-/-^* (n = 5) and *Ifngr1^+/+^* (i.e. C57BL/6J) (n = 5) mice were infected with UgCl223 as described above. Survival was monitored based on aforementioned end-point criteria.

C57BL/6J mice were infected with UgCl223 as described above. At 21 days post-infection, mice were intraperitoneally injected with 10 μg recombinant mouse IFNγ (Biolegend) every 24 hours (n = 4). Sterile saline was used as the vehicle control (n = 5). Lung and brain fungal burden was measured at 35 days post-infection as described above.

### CTLA-4 monoclonal antibody blockade

Mice were intranasally infected with UgCl223 and intraperitoneally injected with 0.5 mg of anti-CTLA-4 monoclonal antibody (9D9; Bio X Cell) starting at 21 days post-infection and every 7 days thereafter. Because CTLA-4 blockade does not result in depletion of CTLA-4+ cells, the efficacy was assessed by flow cytometric analysis via FoxP3+ CD4 T-cell depletion (88, 89) (**Supplementary Figure 8**). An IgG2b (MPC-11; Bio X Cell) isotype antibody was used as a control. Lung and brain fungal burden was measured at 35- and 49-days post-infection as described above. The following fluorophore-labelled antibodies were used for this experiment: CD45 for IV label (30-F11, APC-eFluor780, eBioscience), CD45 (30-F11, BUV805, BD Biosciences), B220 (RA3-6B2, BV650, Biolegend), CD11c (N418, APC-eFluor780, Thermo Fisher Scientific), CD11b (M1/70, APC-eFluor780, Thermo Fisher Scientific), F4/80 (BM8, APC- eFluor780, Thermo Fisher Scientific), TCRβ (H57-597, PE, eBioscience), CD4 (GK1.5, BUV39, BD Biosciences), CD8 (53-6.7, AF700, Thermo Fisher Scientific), CD44 (IM7, PerCp-Cy5.5, Biolegend), PD-1 (J43, APC, Invitrogen), FoxP3 (FJK-16S, AF488, eBioscience), CTLA-4 (UC10-4B9, BV605, Biolegend).

### Statistics

Statistical analysis was performed with GraphPad Prism 9 software (La Jolla, CA). Power calculations were performed to assess appropriate sample size for all experiments. Data were analyzed using one or two-tailed t-test, one-way or two-way ANOVA. Bonferroni adjustment for multiple comparisons was implemented when justified. P-values ≤ 0.05 were considered statistically significant. All data presented in this study, except in **Supplementary Figure 1** and **Supplementary Figure 7-8**, are representative from a minimum of at least three independent experiments or biological replicates.

## ACKNOWLEDGEMENTS

We thank Dr. Marc Jenkins and Jennifer Walter for kindly gifting the Tbet-zsGreen FoxP3-RFP mice used in this study. We also thank the University of Minnesota Flow Cytometry Resource (UFCR) for technical assistance in this study. Sequencing for this study was performed by the Roy J. Carver Biotechnology Center at the University of Illinois at Urbana-Champaign, which was supported in part by the University of Minnesota Genomic Subsidy Program.

Funding was provided by National Institutes of Health grants F30AI155292 to MD, and R01NS412966 and R01AI134636 to KN. In addition, M.D. was supported by the University of Minnesota Medical Scientist Training Program (MSTP) under NIH grant T32GM008244 and University of Minnesota Lung Biology Dinnaken Fellowship.

MD and KN conceived the study. MD planned and performed all experiments, except the *Ifngr1^-/-^* survival experiment which was performed by ED. JMY provided technical assistance for the experiments. JD analyzed the single cell RNA sequencing data, with input from MD and KN. MD analyzed the remaining data under the direction of KN. MD wrote the initial draft of the manuscript. MD and KN reviewed and edited subsequent manuscript drafts. All authors have declared that no competing interests exist. All authors have approved the final version of the submitted article.

## DATA AVAILABILITY STATEMENT

The data underlying Figs. 1-5, 8, Supplementary Figs. 1-6, and Supplementary Table 1 are openly available in NCBI SRA under BioProject PRJNA1271018. The accompanying downstream analysis is available in Dryad: https://doi.org/10.5061/dryad.j3tx95xsx. The data underlying Figs. 6-7, 9, and Supplementary Figs. 7-8 are available upon request.

---

**Supplementary Figure 1: Single-cell RNA sequencing analysis of pulmonary CD45 T-cells during *C. neoformans* infection.** C57BL/6J mice were infected with *C. neoformans* UgCl223 at 14 days post-infection (Latent 14 DPI) or 150 days post-infection (Latent 180 DPI), KN99*α* at 14 days post-infection (Lethal 14 DPI), or uninfected. CD45+ cells were isolated from the lung homogenate and processed for single-cell RNA sequencing analysis via the 10x Genomics platform. (A) Heatmap showing SingleR annotation against Seurat clusters. (B) UMAP projection showing Seurat clusters annotated by cell type. (C) UMAP showing only the T-cell cluster in the various experimental conditions. (D) Feature plot showing *Cd4* expression on T-cell cluster in the various experimental conditions. (E) Volcano plot showing differential expression analysis of *Cd4*-expressing cells in the uninfected condition compared to that of all the other experimental conditions (Lethal 14 DPI, Latent 14 DPI, and Latent 150 DPI).

**Supplementary Figure 2: Heatmap of top 10 expressed genes per Seurat cluster of the pulmonary CD4 T-cell response to *C. neoformans* infection.** Heatmap of top 10 genes expressed by the pulmonary CD4 T-cell clusters during *C. neoformans* infection.

**Supplementary Figure 3: Heatmap of top 10 expressed genes per Seurat cluster of the pulmonary effector CD4 T-cell response to *C. neoformans* infection.** Heatmap of top 10 genes expressed by the pulmonary effector CD4 T-cell (cluster 2) subclusters during *C. neoformans* infection.

**Supplementary Figure 4: Th17-related genes are decreased over the course of latent *C. neoformans* infection.** Differential expression analysis of pulmonary effector CD4 T-cell subclusters 1-12 was performed between two experimental conditions: (A) *C. neoformans* KN99*α* at 14 days post-infection (Lethal 14 DPI) and UgCl223 at 14 days post-infection (Latent 14 DPI), and (B) *C. neoformans* UgCl223 at 180 days post-infection (Latent 180 DPI) and Latent 14 DPI. For each comparison, significant Th17-associated genes within each subcluster (with globally adjusted p-value < 0.05) were plotted according to logFC. Positive logFC values indicate higher expression in the first condition listed, whereas negative logFC values indicate higher expression in the second condition. The dashed horizontal line denotes biological significance at |logFC| ≥ 1.5.

**Supplementary Figure 5: *Rorc* (RORγt) and *Bcl6* (Bcl6) expression by pulmonary effector CD4 T-cells is not significantly changed during *C. neoformans* infection.** Feature plots showing expression of (A) *Rorc* (RORγt) and (B) *Bcl6* (BCL6) by pulmonary effector CD4 T-cells across the three experimental conditions: *C. neoformans* KN99*α* at 14 days post-infection (Lethal 14 DPI), UgCl223 at 14 days post-infection (Latent 14 DPI), and UgCl223 at 180 days post-infection (Latent 180 DPI). Cells expressing > 0 copies of the respective gene of interest were quantified and divided by the total number of effector pulmonary CD4 T-cells to determine the proportion of effector CD4 T-cells expressing the respective gene of interest in each biological replicate. Significance was determined via one-way ANOVA with Bonferroni correction.

**Supplementary Figure 6: *Ctla4* (CTLA-4) expression by pulmonary effector CD4 T-cells is significantly increased during *C. neoformans* infection.** Feature plots showing expression of *Ctla4* (CTLA-4), *Pdcd1* (PD-1), *Lag3* (LAG-3), *Havcr2* (TIM-3) by pulmonary effector CD4 T-cells across the three experimental conditions: *C. neoformans* KN99*α* at 14 days post-infection (Lethal 14 DPI), UgCl223 at 14 days post-infection (Latent 14 DPI), and UgCl223 at 180 days post-infection (Latent 180 DPI). Cells expressing > 0 copies of the respective gene of interest were then quantified and divided by the total number of effector pulmonary CD4 T-cells to determine the proportion of effector CD4 T-cells expressing the respective gene of interest in each biological replicate. Significance was determined via two-way ANOVA with Bonferroni correction.

**Supplementary Figure 7: Exhaustion marker flow cytometry gating strategy.** Representative flow scatter plots of a single cell suspension isolated from the lungs of a C57BL/6J mouse at 21 days post-infection with *C. neoformans* UgCl223. Fluorescence minus one (FMO) controls were used to determine gating for the BV605-CTLA4, APC-PD1, BV421-LAG3, and BV711-TIM3 populations.

**Supplementary Figure 8: CTLA4 monoclonal antibody depletes FoxP3+ regulatory CD4 T-cells.** C57BL/6J mice were latently infected with *C. neoformans* UgCl223. At 21 days post-infection (DPI), mice were injected with 0.5 mg anti-CTLA4 monoclonal antibody (9D9) or an IgG2b isotype control via weekly intraperitoneal injections. At 28-, 35-, 42- and 49- days post infection, one representative mouse was injected with CD45-APC-eFluor780 via tail vein and euthanized after 3 minutes. Following PMA/ionomycin stimulation, lungs were perfused with cold PBS and digested via collagenase A. Single cell suspensions were stained and quantified using flow cytometry. Live+, Singlet+, Lung+, B220-, NK1.1-, CD11c-, CD11b-, F4/80-, CD8-, CD4+, CD44+, FoxP3+ cells were visualized and quantified by scatterplot.

**Supplementary Table 1: Standard error of average percentage of pulmonary effector CD4 T-cells expressing lineage-defining transcription factor(s).**

